# Primary Human Cell-Derived Extracellular Matrix from Decellularized Fibroblast Microtissues with Tissue-Dependent Composition and Microstructure

**DOI:** 10.1101/2023.08.15.553420

**Authors:** Vera C. Fonseca, Vivian Van, Blanche C. Ip

**Affiliations:** Department of Pathology & Laboratory Medicine, Brown University. Box G-E5. Providence, RI 02912

## Abstract

Human extracellular matrix (ECM) exhibits complex protein composition and architecture depending on tissue and disease state, which remains challenging to reverse engineer. One promising approach is based on cell-secreted ECM from human fibroblasts, which can then be decellularized into an acellular biomaterial. However, fibroblasts initially seeded on rigid tissue culture plastic or biomaterial scaffolds experience aberrant mechanical cues that influence ECM deposition. Here, we show that engineered microtissues of primary human fibroblasts seeded in low-adhesion microwells can be decellularized to produce human, tissue-specific ECM. We investigate: 1) cardiac fibroblasts, as well as 2) lung fibroblasts from healthy, idiopathic fibrosis and chronic obstructive pulmonary disease donors. We demonstrate optimized culture and decellularization conditions, then characterize gene expression and protein composition. We further characterize ECM microstructure and mechanical properties. We envision that this method could be utilized for biomanufacturing of patient and tissue-specific ECM for organoid drug screening as well as implantable scaffolds.

**Impact:** In this study, we demonstrate a method for preparing decellularized matrix using primary human fibroblasts with tissue and disease-specific features. We aggregate single cell dispersions into engineered tissues using low adhesion microwells and show culture conditions that promote ECM deposition. We demonstrate this approach for cardiac fibroblasts as well as lung fibroblasts (both normal and diseased). We systematically investigate tissue morphology, matrix architecture, and mechanical properties, along with transcriptomic and proteomic analysis. This approach should be widely applicable for generating personalized ECM with features of patient tissues and disease state, relevant for culturing patient cells ex vivo as well as implantation for therapeutic treatments.

## Introduction

Human cells secrete extracellular matrix (ECM) proteins when cultured ex vivo, representing a promising source of human ECM that recapitulates some of the complex bioactivity observed in vivo [1]. However, the composition and microstructure of cell-derived ECM can depend sensitively on the initial culture conditions, whether cells are seeded on 2D tissue culture plastic [2–6] or within 3D biomaterial scaffolds [7–12]. Indeed, dispersed cells cultured on 2D substrates may alter their polarity and differentiated phenotype relative to more cell-dense 3D architectures, which likely affects their secreted ECM [13]. Further, cells cultured on stiffer substrates can exhibit a more inflammatory phenotype associated with fibrosis [14], which may limit the utility of the resulting ECM for tissue regeneration. Nevertheless, the ECM secreted by human cells in the absence of any initial biomaterial cues has not been extensively characterized.

Multicellular microtissues are prepared by aggregating single cell dispersions under low adhesion conditions, representing a facile approach to prepare cell-dense tissues [15]. For instance, multicellular microtissues can be prepared using hanging drops, spinner flasks, or pellet culture. Alternatively, arrays of concave microwells made with non-cell-adhesive hydrogels permit controlled microtissue formation of consistent size and shape. This approach was utilized elsewhere to prepare microtissues of mesenchymal stem cells [16, 17], which were subsequently decellularized as tissue engineering scaffolds.

ECM composition can also depend sensitively on cell type, which has been previously explored using stem cells, chondrocytes, and hepatocytes on tissue culture plastic [1]. ECM deposition from fibroblasts is intriguing due to their functional role in tissue development and regeneration, which can vary across different organs [18]. For example, healthy heart tissue ECM includes collagens, fibronectin, proteoglycans, glycoproteins as well as laminin and elastin [19]. In comparison, healthy lung tissue ECM also includes collagens, fibronectin, glycoproteins as well as elastin [20]. ECM remodeling occurs during chronic lung diseases such as idiopathic pulmonary fibrosis (IPF) or chronic obstructive pulmonary disease (COPD), with increased ECM crosslinking and deposition. Notably, breast cancer cells show distinct morphology and migration when cultured on decellularized matrix secreted by embryonic or tumor associated fibroblasts, respectively [21].

In this article, we decellularize microtissues of human fibroblasts and elucidate tissue-specific differences in the resulting ECM. First, we optimize culture and decellularization conditions for primary cardiac fibroblasts, which are characterized using histology and proteomics. We demonstrate that decellularized matrix secreted by cardiac fibroblasts can support the proliferation of human cardiac myocytes and microvascular endothelial cells. Second, we investigate the secreted matrix from primary lung fibroblasts from healthy donors and IPF patients, revealing pronounced microstructural differences based, as well as differences in proteomic composition. Finally, we compare the mechanical properties and gene expression profiles across cardiac and lung derived microtissues. Overall, we demonstrate the potential of this method for recapitulating the composition and architecture of healthy and diseased extracellular matrix in a tissue-specific manner.

## Methods

### Cell culture

Human cardiac fibroblasts (C-12375, PromoCell) were expanded in Fibroblast Growth Medium 3 (C-23025, PromoCell). Human cardiac myocytes (HCM; C-12810, PromoCell) were expanded in Myocyte Growth Medium 3 (C-22070, PromoCell). Human cardiac microvascular endothelial cells (C-12285, PromoCell) were expanded in Endothelial Cell Growth Medium MV (C-22020, PromoCell). Human lung fibroblasts (CRL-4058, ATCC), were expanded in Fibroblast Basal Medium (PCS-201-030, ATCC) supplemented with Fibroblast Growth Kit-Low Serum (PCS-201-041, ATCC) and puromycin (Gibco A1113803) at a concentration of 0.3 µg/mL. Human lung fibroblasts from IPF (PCS-201-020, ATCC) or COPD (PCS-201-017, ATCC) patients were expanded in Fibroblast Basal Medium supplemented with Fibroblast Growth Kit-Low Serum. Fibroblasts were treated with or without recombinant human transforming growth factor beta 1 (TGF-β1) protein (240-B, R&D Systems) at 1.25 ng/mL. Cells were trypsinized, counted, and re-suspended to the desired cell density for each experiment.

MDA-MB-231 breast adenocarcinoma cell lines stably transduced with green fluorescent protein fused to nuclear histones (H2B-GFP) were a gift from R. Weissleder (Massachusetts General Hospital). Cells were maintained in DMEM with L-Glutamine, 4.5 g/L glucose and sodium pyruvate (Gibco 11965092), with 10% fetal bovine serum (Gibco 16000044) and 1% penicillin-streptomycin (Gibco 15140122). Cells were regularly subcultured at 1:10 split below passage number 25.

### Micro-mold fabrication, and formation of microtissues

2% w/v agarose micro-wells were prepared as described [22]. Agarose gels were made with powdered agarose (BP160, ThermoFisher), autoclaved, then dissolved in sterile phosphate buffered saline (PBS; HyClone SH30256.FS, ThermoFisher). Gels with round micro-wells (400 to 800 µm in diameter) with 35, 96, 256 or 721 micro-wells per gel were used to aggregate spheroidal microtissues. Agarose molds were placed inside sterile cell culture plates and equilibrated either with Fibroblast Basal Medium (lung) or Fibroblast Growth Medium 3 without serum (cardiac) with three exchanges prior to microtissue fabrication.

Equilibrated agarose gels were seeded with dispersed single cells at a density of 375 to 3,000 cells per spheroid micro-well. Cells were allowed to settle to the bottom of the micro-wells for 30-45 minutes prior to adding complete media (containing serum). Cell seeded gels were cultured for 3, 6, 9 or 12 days in a humidified incubator at 37°C with 5% CO_2_, with media changes every 3 days. Cells aggregated into one spheroid-shaped microtissue per micro-well within 4-24 hours after seeding.

### Morphological Characterization

Microtissues in agarose gels were imaged with an inverted microscope (Zeiss Axio Observer Z1). The cross-sectional area of microtissues were measured using ImageJ.

### Decellularization of microtissues

We optimized our decellularization protocol based on previously published methods to successfully remove all cellular material and retain much of the ECM [23–27]. Microtissues in agarose gels were washed with PBS three times. Decellularization of microtissues were either completed with microtissues remaining in the agarose gels, or after microtissues were collected into a tube. Microtissues in gels or in tubes were first treated with three rounds of 0.5% Triton-X100 (T9284, MilliporeSigma) in 20 mM NH4OH (09859, MilliporeSigma) in sterile PBS with protease inhibitors (PI; PI78439, ThermoFisher) for 45 mins, followed by three rounds of washes with sterile PBS + PI for 45 mins, then subjected to one round of incubation with DNase I (4716728001, MilliporeSigma) + RNase A (19101, Qiagen) + PI for 72 hours, followed by three rounds of washes with sterile PBS + PI for 45 mins. All steps were done at 37°C with agitation. The resulting cultured ECM were immediately used for mechanical testing with atomic force microscopy (AFM), stored in sterile PBS at 4°C for visual inspection, fixed in 10% buffered formalin for histological analysis, or snap-frozen in liquid nitrogen and stored at −80°C for subsequent biochemical analysis.

### Histology

Non-decellularized or decellularized microtissues cultured for different days were fixed in 10% buffered formalin (427098, ThermoFisher) in the agarose gels, paraffin-embedded, sectioned at 5 µm then stained with hematoxylin and eosin (H&E) or Sirius Red^TM^ (24901-250, Polyscience) to examine microtissue morphology or fibrillar collagen deposition, respectively.

### Second harmonic generation microscopy of fibrillar architecture

Microtissues cultured for 6 days were fixed in 10% buffered formalin in the agarose gels. Fibrillar collagen was visualized using an Olympus FV-1000-MPE multiphoton equipped with a Mai Tai HP tunable laser with the excitation wavelength set to 790 nm and a 405/40 filter cube to select for fibrillar collagen second-harmonic signal. 10% formalin was replaced with FBS prior to imaging 25x (NA 1.05, WD 2 mm) dipping objective.

### Proteomic Analysis

Microtissues in agarose gels were washed with PBS three times then collected into a tube. ECM-enrichment and proteomics procedures were adapted from Naba et. al. [28]. The Millipore Compartment Protein Extraction Kit (2145, MilliporeSigma) was used to concentrate ECM proteins for proteomics analysis of the microtissues. LC-MS/MS identification and label free quantitation of ECM proteins were conducted by The University of Oklahoma Proteomics Core (Supplementary Methods). Proteome Discoverer 2.3 (1% FDR) were used for the proteomics analysis of ECM proteins. The open-access MatrisomeDB resource [29] was used to identify ECM proteins, then classified them into six categories: collagens, ECM regulators, ECM-affiliated proteins, glycoproteins, proteoglycans, and secreted factors. An established iBAQ algorithm as described by Schwanhausser et al. [30] was used to semi-quantity ECM components (by % molar of total ECM proteins) by dividing each individual protein’s total intensity with the theoretical number of tryptic peptides between 6-30 amino acids in length (PeptideMass, SIB Swiss Institute of Bioinformatics).

### dsDNA, collagen and sulphated glycosaminoglycans (sGAG) concentration measurements

DsDNA concentration of decellularized ECM were measured with QUANT-IT™ PICOGREEN™ dsDNA Assay Kit (P7589, ThermoFisher) per manufacturer protocol [31]. Collagen content of flash-frozen microtissues was measured using a modified hydroxyproline assay [32]. sGAG content of flash-frozen microtissues was measured using the 1,9-dimethylmethylene blue (DMMB) assay [33, 34]. Decellularized or flash-frozen microtissues were digested in papain solution (125 µg/mL, P4762, MilliporeSigma) for 72 hours at 65°C prior to measurements with corresponding methods.

### AFM of microtissue mechanical properties

Samples tested went through a decellularization process, were plated on collagen-coated coverslips and incubated on coverslips at 4°C for 48 hours prior to testing. Force measurements were collected using an atomic force microscope (Asylum MFP-3D-BIO) connected to a Nikon Eclipse Ti-U epifluorescence microscope. Multiple testing sessions were conducted for the various samples to account for systematic errors.

All experiments were carried out at room temperature in fluid environments. The AFM was allowed to equilibrate before tests to minimize deflection laser and/or piezo drift. Force maps were collected for a variety of samples using a force mapping technique in contact mode. Briefly, individual force curves were taken at discrete points across a region of interest. During analysis, the spatial arrangement of the data was retained to create a matrix of elastic modulus values. Force–indentation data were sampled using a cantilever with spring constant of 0.03 N/m at 5 kHz with an approach velocity of 10 μm/sec. A trigger force of about 4 nN was used for all samples with the deflection set to 100 nM. Scan size used was 5 μm and the resolution was 4 × 4 pts. Force versus indentation data were analyzed using custom MATLAB scripts (MathWorks) utilizing the Hertz contact model.

### RNA-sequencing (RNAseq)

Total RNA was extracted from snap-frozen microtissues using Quick-RNA Miniprep Kit (R1054, Zymo Research) per manufacturer protocol. RNA concentration was measured using NanoDrop (ThermoFisher), and qualified using the Bioanalyzer (Agilent), where all samples had RNA integrity number (RIN) > 9.0. Genewiz performed the RNA library preparation and RNAseq using the Illumina HiSeq®.

### Cell adhesion to decellularized ECM

Decellularized agarose gels with or without adult heart microtissues were seeded with myocytes or endothelial cells and cultured for 24 hours. To examine proliferation, the nucleoside analog EdU (5-ethynyl-2’-deoxyuridine) was then added. Cells on ECM in EdU were cultured for another 24 hours (HCMEC) or 48 hours (HCM), then fixed in 10% formalin for immunohistochemical evaluation using the Click-iT™ EdU Cell Proliferation Kit (C10634, ThermoFisher) per manufacturer’s protocol.

Decellularized agarose gels with or without adult healthy or IPF lung microtissues were seeded with MDA-MB-231 and incubated for 10 days. Agarose gels were imaged with a widefield inverted fluorescent microscopy fitted with camera (Zeiss Axio Observer Z1), using a 10X objective and filter set with excitation (470/40 nm) and emission cutoffs (525/50 nm) for GFP at the same exposure time. Fluorescent intensity was measured with ImageJ to quantify proliferation of cells.

### Statistical analysis

JMP® Pro 17 software (SAS Institute) was used for statistical analyses. Data are presented as mean ± SD. A minimum of N = 3 per group for all experiments. All data were first examined for equal variance and for normality using the Shapiro–Wilk test. Student’s two-sided *t*-test or one-way ANOVA with post-hoc Tukey HSD Test, or their non-parametric equivalents were used. Statistical significance was set at P ≤ 0.05.

## Experiment

### Cardiac Fibroblasts Form Multicellular Microtissues and Deposit Fibrillar Collagen

Single cell dispersions of cardiac fibroblasts were seeded in agarose microwells at densities ranging from 375-3000 cells per well. After 1 day, these cells were assembled into multicellular microtissues with cross-sectional area (11,065 to 41,550 μm^2^) increasing with seeding densities (**Figure 1AB**). After 6 days, larger (1500, 3000 cells/well) but not smaller microtissues compacted (**Figure 1AB**). Hematoxylin and eosin (H&E) staining of day 6 microtissue sections showed an asymmetric distribution of cells (**Figure 1C**), and no evidence of a necrotic core for all microtissue sizes, suggesting that oxygen and nutrient transport via diffusion was adequate. Sirius red staining revealed patches of fibrillar collagen within the microtissue, occupying about half of this cross-sectional area (**Figure 1D**). The dotted circles in **Figure 1CD** suggested the exclusion of cells in collagen-rich regions.

**Figure 1.**
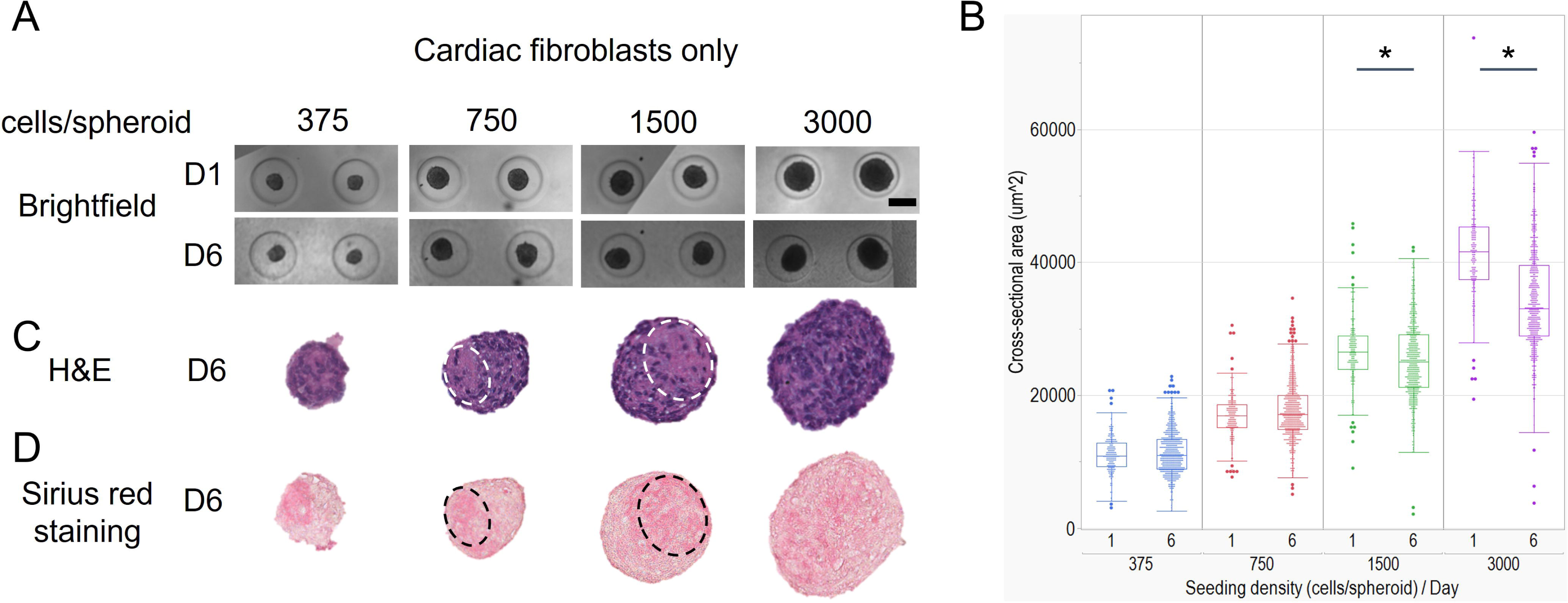
Human cardiac fibroblasts (HCF) form multicellular microtissues and deposit fibrillar collagen. Size of microtissues are depending on seeding density of HCF cultured in low adhesion agarose microwells (A). Larger microtissues (1500 or 3000 cells/well) but not smaller microtissues compacted from day 1 to day 6 (B). Hematoxylin and eosin (H&E) staining of day 6 microtissue sections showed an asymmetric distribution of cells without necrotic core (C). Sirius red stain reveals fibrillar collagen deposited by HCF, and the dotted circles suggested the exclusion of cells in collagen rich regions. Student’s t-test or the non-parametric equivalent was used to examine statistical significance between maturation days. * denotes statistical significance (p-value ≤ 0.05).

The kinetics of collagen deposition were then quantified by characterizing multicellular microtissues (1000 cells per well) over a 12-day period. Multicellular microtissues exhibited a cross-sectional area of 24,001 μm^2^ (**Figure 2AB**) on day 3, and reduced slightly to 22,686, 23,009 and 21,610 μm^2^on day 6, 9 and 12, respectively (**Figure 2AB**). The cells remained uniformly dispersed on day 3 (**Figure 2C**), with only a few small patches of fibrillar collagen evident (**Figure 2D**). Cells became asymmetrically sorted to one side of the microtissue from day 6 onwards, with a concentrated patch of fibrillar collagen on the opposite side (**Figure 2CD**). Thus, 6 days of culture were sufficient for substantial collagen deposition within these cardiac fibroblast microtissues.

**Figure 2.**
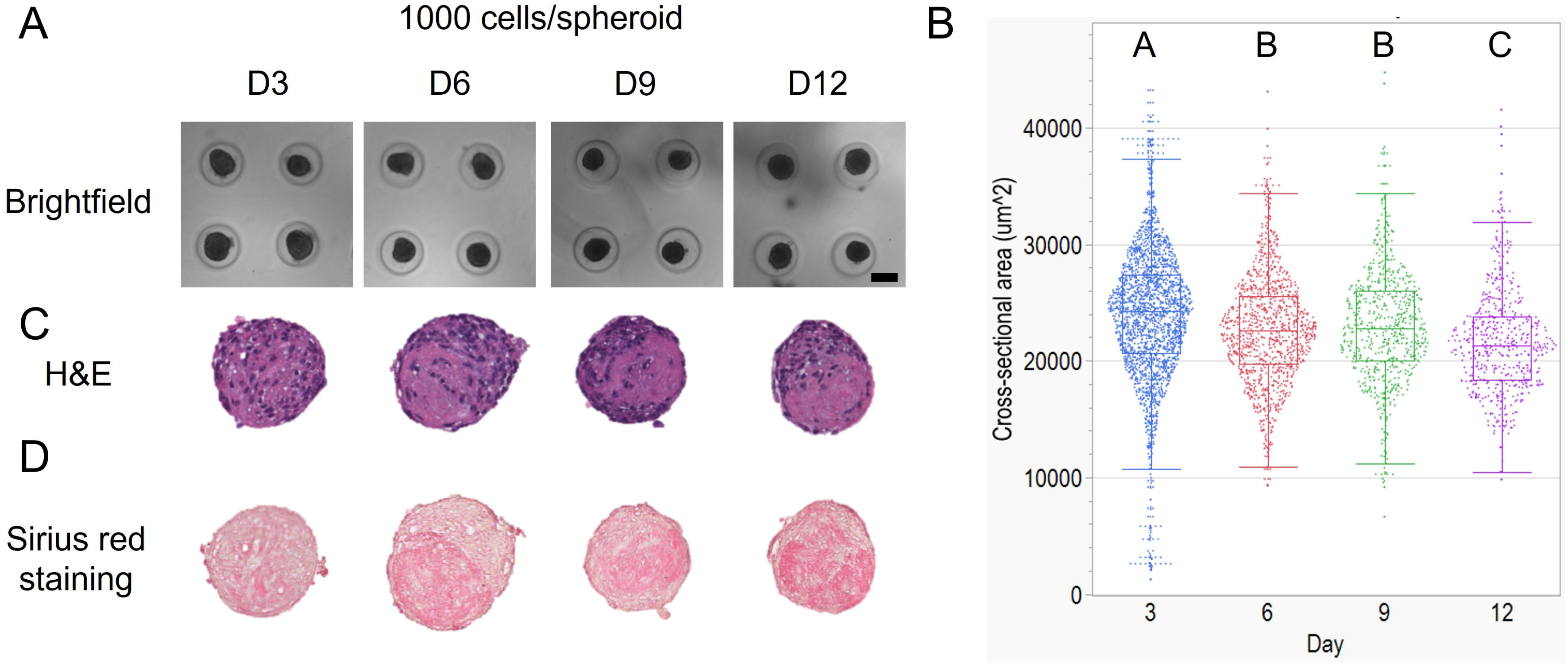
Kinetics of microtissue maturation and collagen deposition by human cardiac fibroblasts (HCF). HCF microtissues with 1000 cells compacted over the course of 12 days (A), with mean cross-sectional areas of 24,001, 22,686, 23,009 and 21,610 μm2 on day 3, 6, 9 and 12, respectively (B). Hematoxylin and eosin (H&E) staining of microtissue sections showed an asymmetric distribution of cells without necrotic core at all four time points (C). Fibrillar collagen deposited by day 3, and increased substantially by day 6 and onwards (D).

For comparison, a tri-culture (HCF_M) of cardiac fibroblasts, myocytes, and endothelial cells at a ratio of 6:2:2 that mimic the human myocardium [35, 36] were seeded at 750 cells per well for 6 days (**Figure 3A**). These optimized experimental parameters were chosen based on the previous results with cardiac fibroblasts only (HCF) with no compaction and sufficient collagen deposition. H&E staining shows similar microtissue morphology between HCF and HCF_M microtissues (**Figure 3B**). Moreover, Sirius red staining of HCF_M shows smaller patches of fibrillar collagen within the microtissue relative to HCF, occupying approximately a quarter of the cross-sectional area (**Figure 3C**).

**Figure 3.**
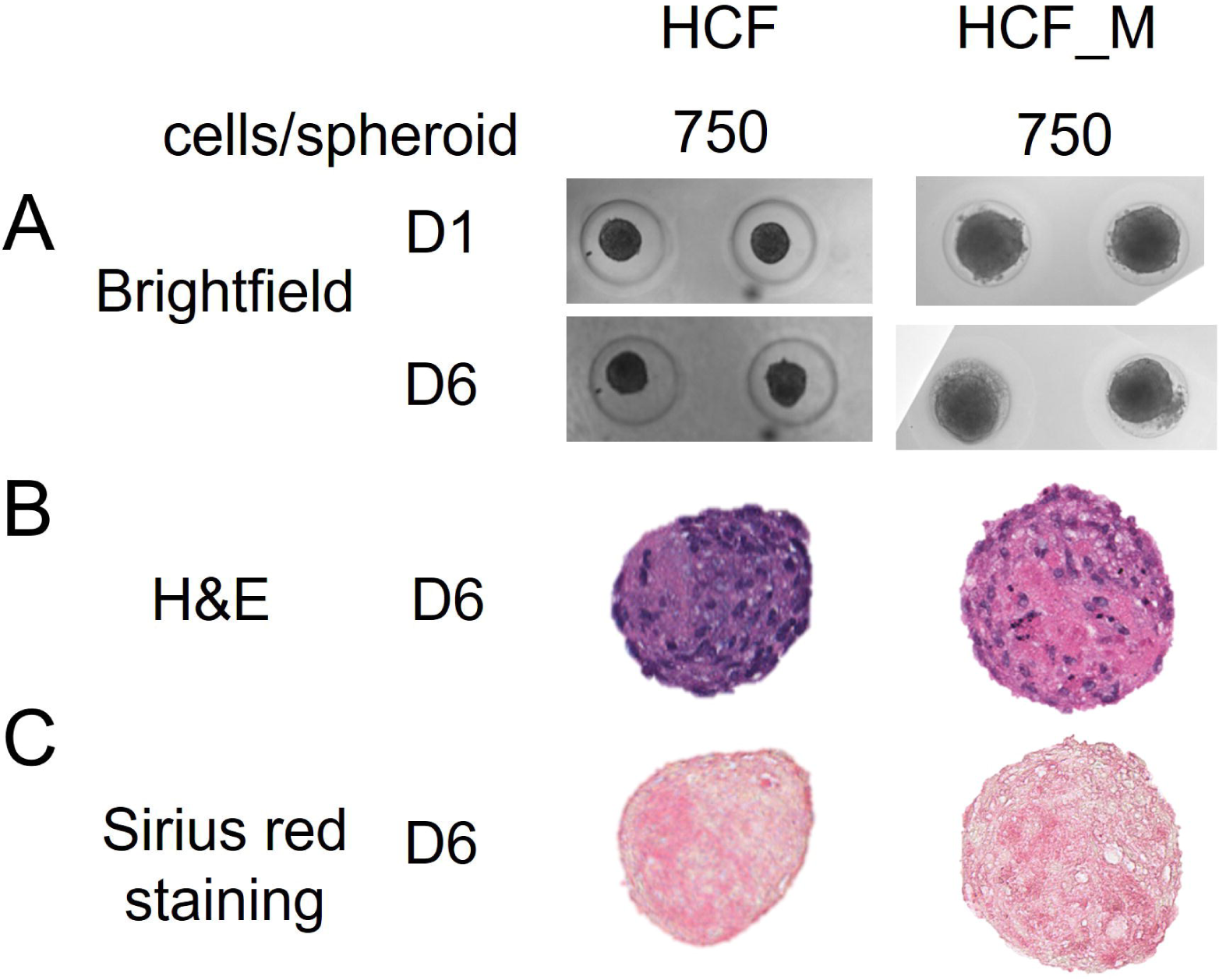
Tri-culture of human cardiac fibroblasts (HCF), cardiac myocytes and cardiac microvascular endothelial cells formed stable microtissues and deposited fibrillar collagen. Tri-culture microtissues (HCF_M) appeared less compacted than HCF microtissues (A). Hematoxylin and eosin (H&E) staining of microtissue sections showed an asymmetric distribution of cells without necrotic core for both HCF and HCF_M (B). Both HCF and HCF_M deposited similar patterns of fibrillar collagen as shown by Sirius red staining (C).

Proteomic analysis of cardiac microtissues revealed ECM protein composition consistent with those previously identified in human and porcine cardiac tissues [37, 38], showing that HCF and HCF_M shared relatively similar proteomes. HCF_M had significantly greater amounts of ECM-affiliated proteins, but lesser amounts of glycoproteins and proteoglycans, as compared to HCF (**Table 1**). Differentially secreted ECM proteins are listed in **Supplementary Table 1**. We further quantified collagen and sGAG content of Day 6 HCF using a modified hydroxyproline assay and the DMMB assay, respectively. We found that HCF deposited 2.2 ± 0.17 μg (per 10^6^ cells) of collagen and 6.16 ± 0.42 μg (per 10^6^ cells) of sGAG. Since the ECM composition is comparable between HCF and HCF_M, it is likely that most of the ECM is deposited by cardiac fibroblasts within the tri-culture at this cell ratio.

**Table 1:** Proteomics analysis of human cardiac microtissues made with human cardiac fibroblasts only (HCF) and tri-culture of fibroblasts, cardiac myocytes and cardiac microvascular endothelial cells (HCF_M). Student’s t-test or the non-parametric equivalent was used to examine statistical significance between the two cardiac microtissues for each of the six ECM protein categories. * denotes statistical significance (p-value ≤ 0.05).

|  | Collagen |  | ECM Regulators |  | ECM-affiliated Proteins * |  | Glycoproteins * |  | Proteoglycans * |  | Secreted Factors |  |
| --- | --- | --- | --- | --- | --- | --- | --- | --- | --- | --- | --- | --- |
|  | Mean | SD | Mean | SD | Mean | SD | Mean | SD | Mean | SD | Mean | SD |
| **HCF** | 3.62 | 0.33 | 6.34 | 1.26 | 21.68 | 2.64 | 40.81 | 3.66 | 3.72 | 0.17 | 23.82 | 1.96 |
| **HCF_M** | 2.85 | 0.59 | 6.95 | 0.40 | 26.92 | 0.66 | 35.09 | 0.74 | 2.77 | 0.21 | 25.41 | 1.68 |

### Decellularized Cardiac Microtissues Support Human Cardiac Myocyte and Microvascular Endothelial Cell Culture

Cardiac fibroblast microtissues after 6 days were decellularized either inside their individual agarose microwells (in situ decellularization; **Figure 4A**), or after HCF microtissues were transferred into tubes (**Figure 4B**), to generate cultured cardiac ECM. Complete removal of cells was confirmed visually by H&E staining, with no nuclei visible (**Figure 4C**), and cultured cardiac ECM had less than 50 ng/mg ECM dry weight. Dry weights of separate samples with identical numbers of microtissues were determined after lyophilization for 48 hours (**Figure 4B**). Cultured cardiac ECM retained considerable fibrillar collagen as observed initially in the multicellular microtissue (**Figure 4C**).

**Figure 4.**
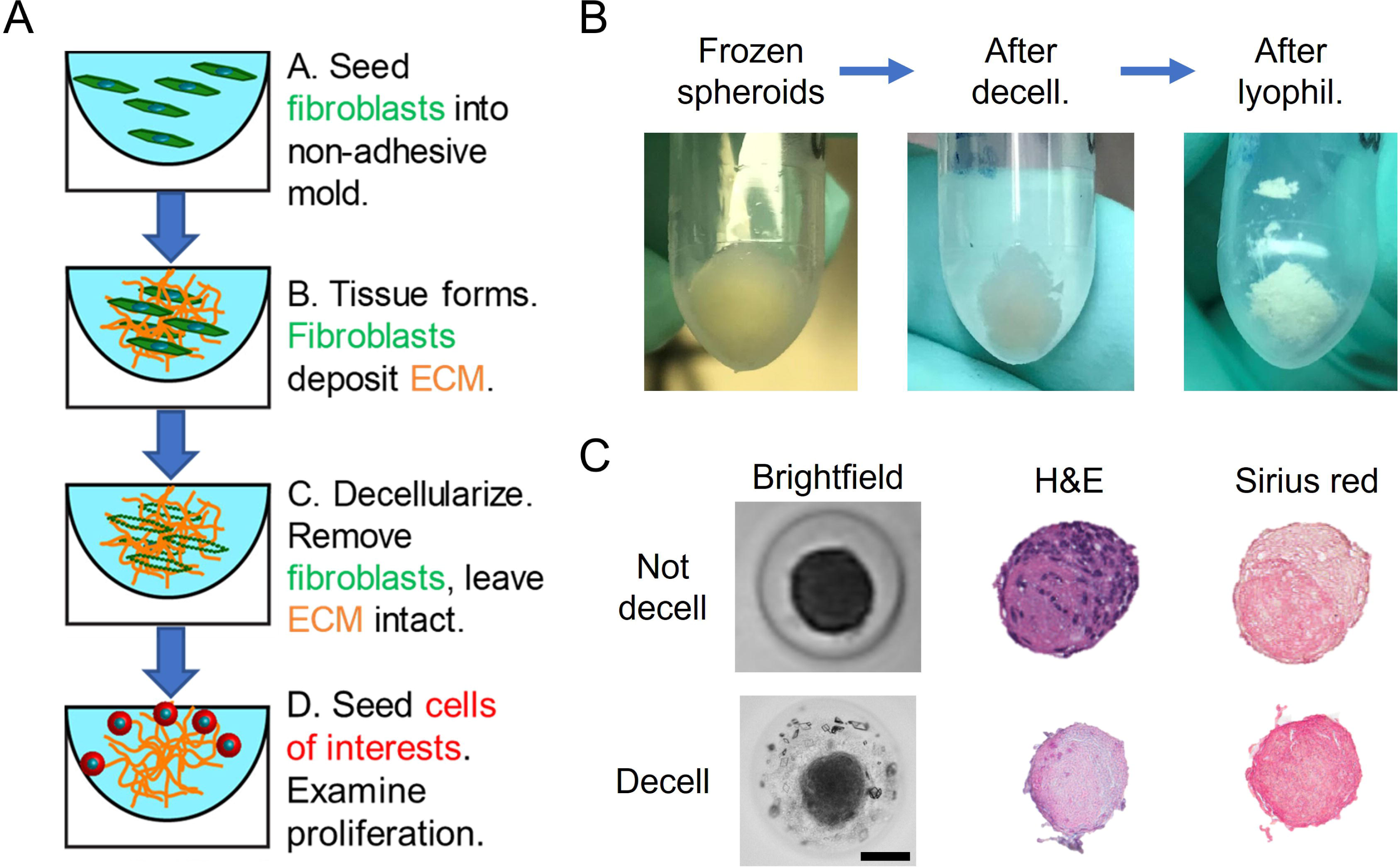
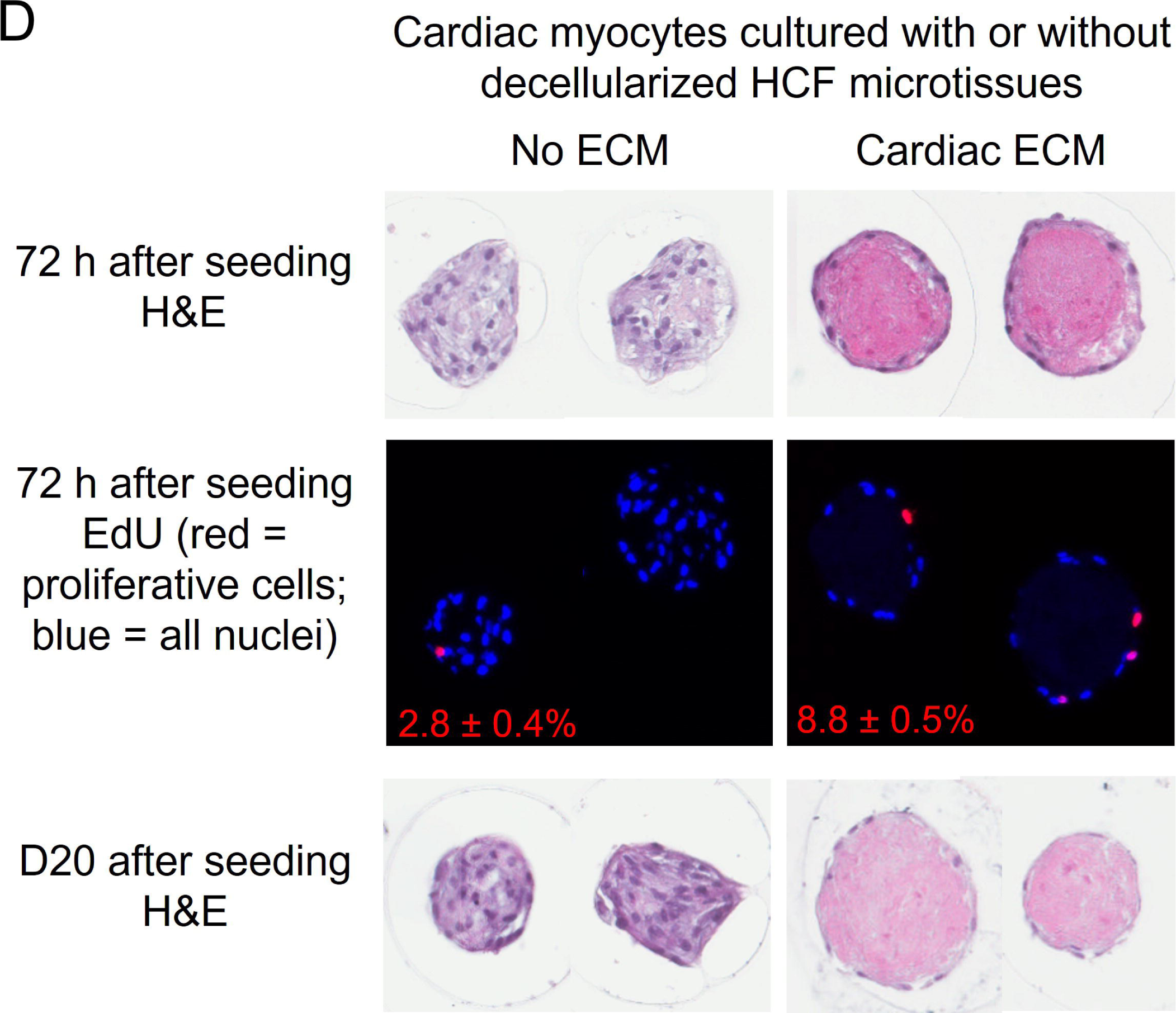
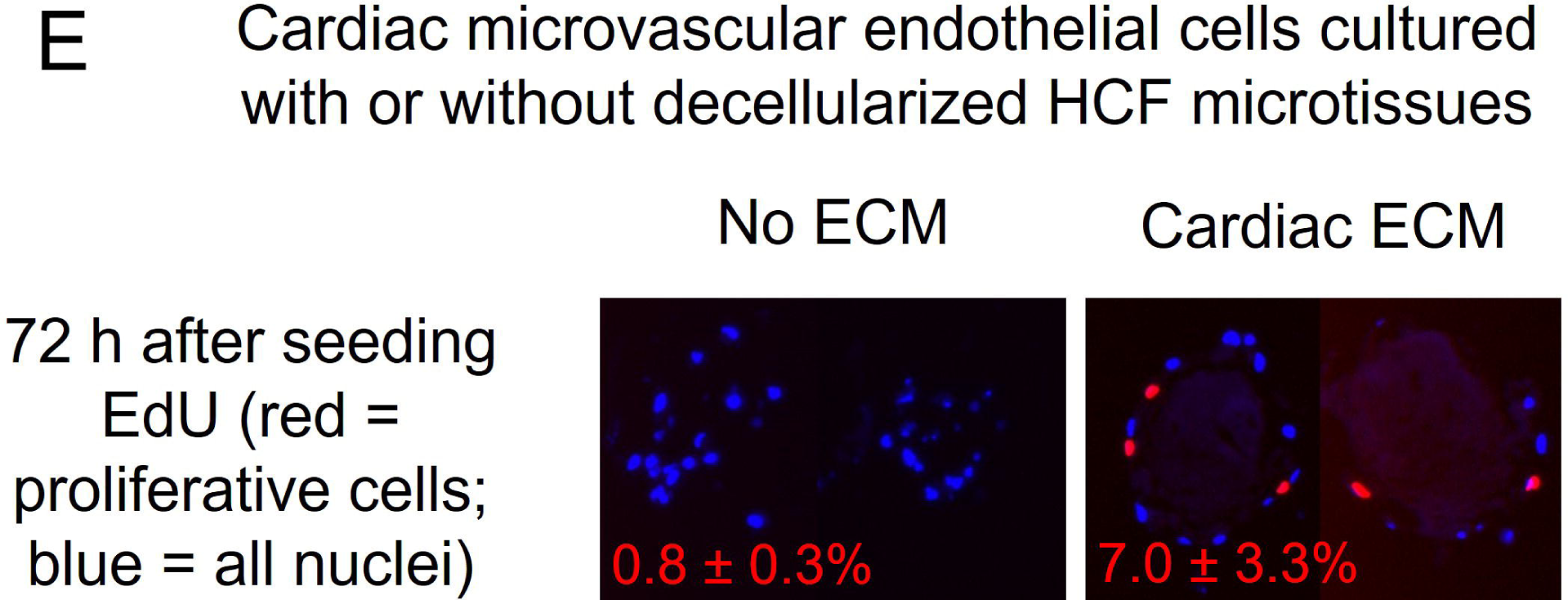
Decellularization of human cardiac fibroblasts microtissues (HCF). HCF after 6 days were decellularized either inside their individual agarose microwells (in situ decellularization; A), or after HCF microtissues were transferred into tubes (B), to generate cultured cardiac ECM. Cultured cardiac ECM were flash-frozen and lyophilized for 48 hours to measure ECM dry weight. Hematoxylin and eosin (H&E) staining of microtissue and cultured cardiac ECM sections showed complete removal of nuclei (C). Sirius red staining revealed that decellularization retained considerable fibrillar collagen (C). Human cardiac myocytes (HCM) were seeded into decellularized agarose gels with or without cultured cardiac ECM and cultured for 72 hours to evaluate proliferation with the Click-iT™ EdU assay, or 20 days to investigate viability. HCM aggregated as multicellular microtissues in the (empty) agarose molds without cultured cardiac ECM, whereas they adhered around the periphery of cultured cardiac ECM (D). HCM had greater proliferation in the presence (8.8% EdU+ nuclei) than in the absence (2.8% EdU+ nuclei) of cultured cardiac ECM (D), and retained good viability out to 20 days after seeding (D). Human cardiac microvascular endothelial cells (HCMEC) had greater proliferation in the presence (7.0% EdU+ nuclei) than in the absence (0.8% EdU+ nuclei) of cultured cardiac ECM (E). Data = mean ± SD.

We investigated whether the cultured cardiac ECM can support the viability and proliferation of human cardiac myocytes (HCM) and cardiac microvascular endothelial cells (HCMEC). HCM were seeded into decellularized agarose gels with or without cultured cardiac ECM and cultured for 72 hours to evaluate proliferation with the Click-iT™ EdU assay, or for 20 days to investigate viability. HCM aggregated as multicellular microtissues in the (empty) agarose molds without cultured cardiac ECM, whereas they adhered around the periphery of cultured cardiac ECM (**Figure 4D**). HCM had greater proliferation in the presence (8.8% EdU+ nuclei) than in the absence (2.8% EdU+ nuclei) of cultured cardiac ECM (**Figure 4D**), and retained good viability out to 20 days after seeding (**Figure 4D**). We demonstrated similar viability and proliferation support of cultured cardiac ECM on HCMEC (**Figure 4E**), where HCMEC-only microtissues exhibited limited proliferation (0.8% EdU+ nuclei), relative to the HCMEC grown on cultured cardiac ECM (7.0% EdU+ nuclei).

### Healthy and Fibrotic Fibroblasts Deposit Different ECM Architecture in Microtissues

As a second case study, multicellular microtissues were prepared using primary human lung fibroblasts, from healthy as well as idiopathic fibrosis (IPF) donors. We hypothesized that IPF lung fibroblasts would exhibit dysregulated ECM deposition relative to healthy fibroblasts, which would be evident in the resulting decellularized microtissue microstructure and composition. Further, exogenous TGF-beta is associated with myofibroblast differentiation, which might further exacerbate aberrant ECM deposition [39].

When seeded in low-adhesion microwells, lung fibroblasts at a seeding density of 800 cells per well typically formed microtissues of ∼175 μm diameter within 24 hours of seeding. Notably, H&E and Sirius red staining showed that healthy lung fibroblast microtissues exhibited relatively sharp collagen bands organized as webs around the nuclei (**Figure 5, 6**). In comparison, healthy lung fibroblast microtissues after TGF-beta treatment exhibited more diffused and less web-like fibrillar collagen (**Figure 5**). IPF lung fibroblast microtissues also exhibited more diffuse collagen than healthy lung fibroblast microtissues, although there was a bit more patchy and nodule-like collagen within certain regions based on second-harmonic generation (SHG) microscopy (**Figure 6B**, orange arrows indicating nodule-like collagen). This is in contrast with the web-like fibrillar collagen in healthy lung microtissues (**Figure 6A**), and the larger but relatively uniform fibrillar collagen region in healthy cardiac microtissues (**Figure 6C**). Lastly, IPF lung fibroblast microtissues after TGF-beta treatment exhibited a decrease in the fibrillar collagen definition relative to untreated IPF lung fibroblast microtissues (**Figure 5**). These trends were corroborated by quantification of collagen and sGAG concentration, which showed a progressive decrease in absolute collagen concentration from 6.4 μg to 2.2 μg and increase in sGAG concentration with TGF-beta treatment and for IPF cells from 2.8 μg to 6.2 μg (**Table 2**).

**Figure 5.**
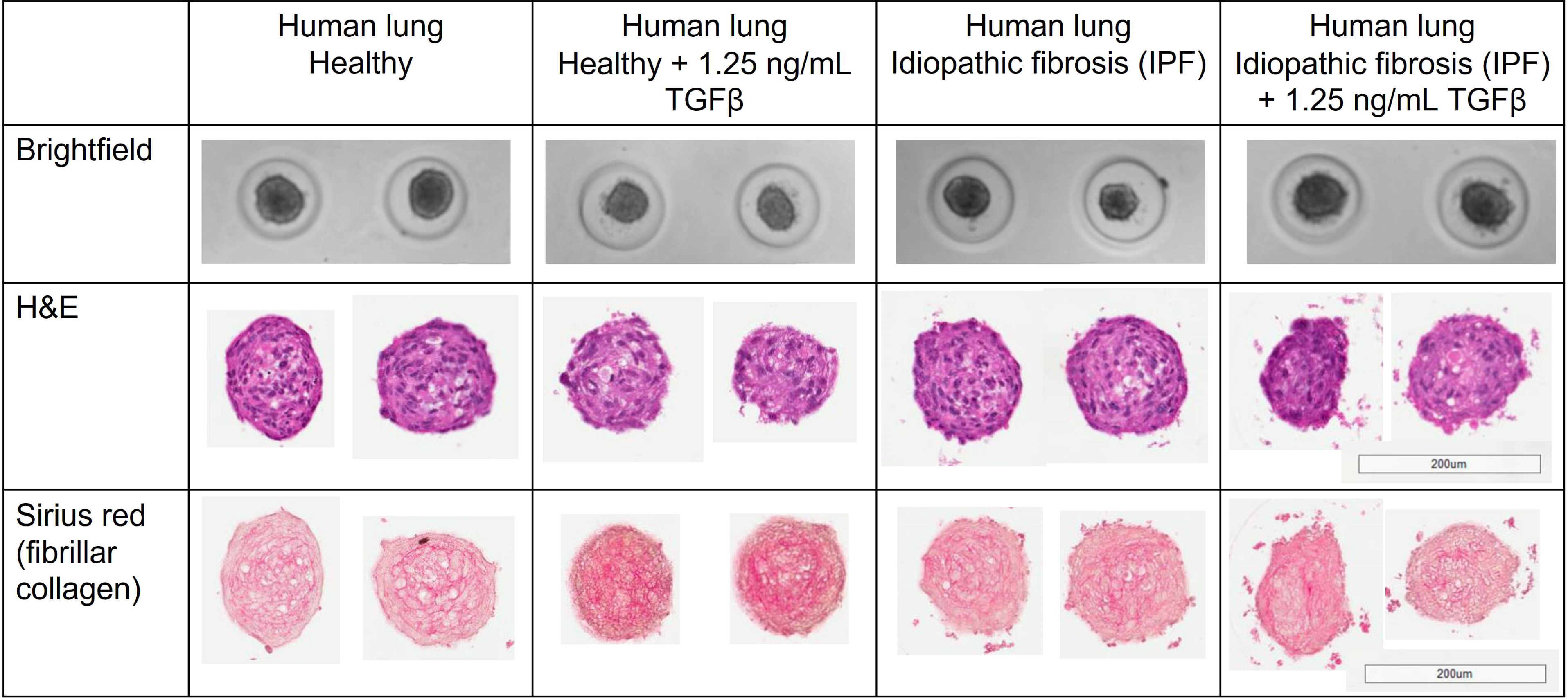
Healthy and fibrotic fibroblast microtissues exhibit differential fibrillar collagen architecture. Representative brightfield images showed that lung fibroblasts isolated from healthy or idiopathic fibrosis donors formed stable microtissues in the presence or absence of TGF-beta. Hematoxylin and eosin (H&E) staining of microtissue sections showed an even distribution of cells without necrotic core for all four microtissues (B). Sirius red staining showed that healthy lung fibroblast microtissues exhibited relatively sharp fibrillar collagen bands organized as webs around the nuclei, whereas microtissues of IPF and in the presence TGF-beta displayed more diffused and disorganized fibrillar collagen.

**Figure 6.**
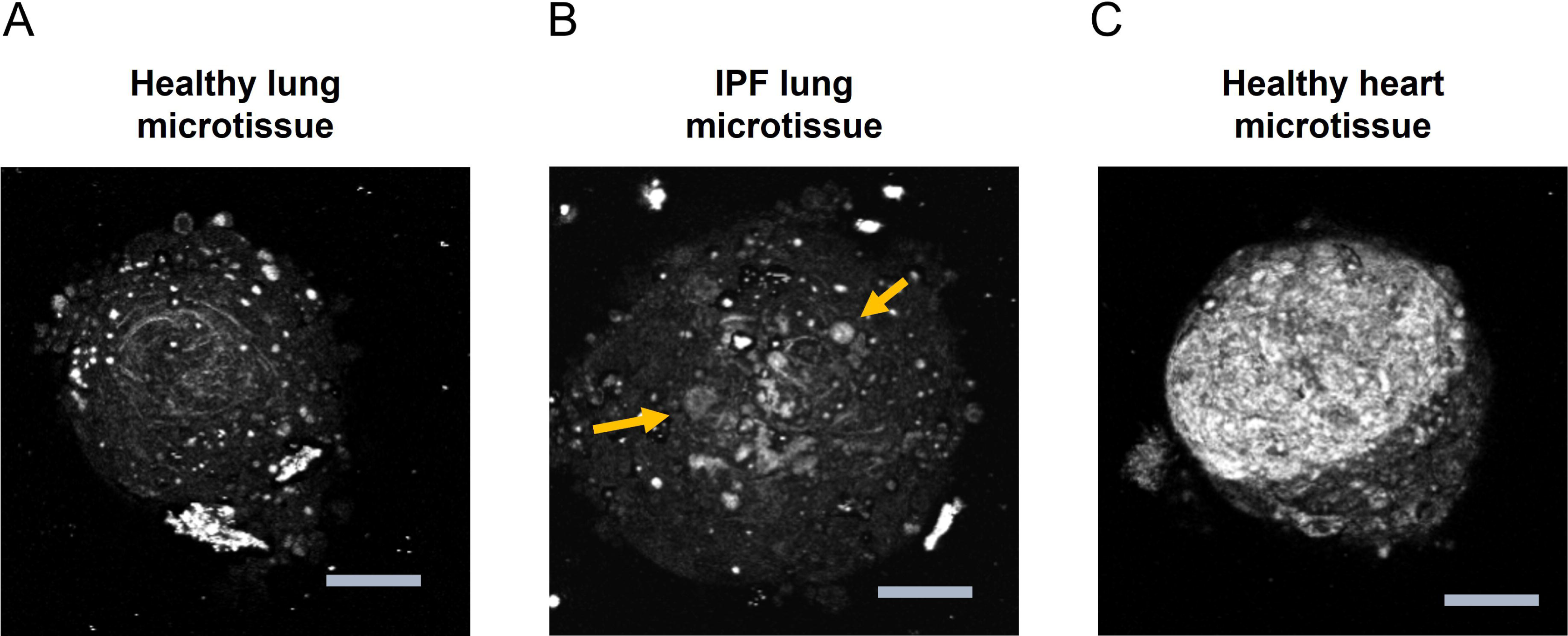
Second-harmonic generation (SHG) microscopy revealed tissue and disease specific collagen architecture of microtissues. Healthy lung microtissues exhibited uniform, web-like collagen architecture (A), whereas microtissues fabricated with lung fibroblasts from idiopathic fibrosis (IPF) donors displayed more localization (patchy) and nodule-like collagen within certain regions (B). Collagen architecture of both lung microtissues are in contrast with the larger but relatively uniform fibrillar collagen region in healthy heart microtissues (C).

**Table 2.** Collagen and sulphated glycosaminoglycans (sGAG) content of four different human fibroblast microtissues. Data = mean ± SD.

| Microtissue | Collagen (μg) | sGAG (μg) |
| --- | --- | --- |
| Healthy Adult Lung | 6.35 ± 2.42 | 2.83 ± 0.50 |
| Healthy Adult Lung + TGF-beta | 4.84 ± 0.82 | 4.81 ± 0.09 |
| Fibrotic Adult Lung | 2.87 ± 0.51 | 5.11 ± 1.68 |
| Healthy Adult Heart | 2.20 ± 0.17 | 6.16 ± 0.42 |

Proteomic analysis also showed a progressive decrease in relative collagen composition, from 28% in healthy lung down to 14% in healthy lung + TGF-beta, then further down to 8.9% for fibrotic lung (**Table 3**). These proteomics results correspond to the measured absolute collagen concentration with hydroxyproline assay (**Table 2**). Addition of TGF-beta to IPF lung fibroblasts did not alter the % collagen in the proteomics samples (**Table 3**). Similar to IPF, microtissues made with lung fibroblasts isolated from COPD patients had less % collagen than healthy lung microtissues. In contrast, secreted factors were significantly elevated in IPF, IPF + TGF-beta, and COPD microtissues as compared to healthy lung microtissues. There is a trend but non-significant increase in ECM-affiliated proteins in IPF, IPF + TGF-beta, and COPD microtissues as compared to healthy lung microtissues. Glycoproteins were typically enriched after TGF-beta treatment relative to untreated, for both normal (significant) and IPF microtissues (non-significant).

**Table 3.**
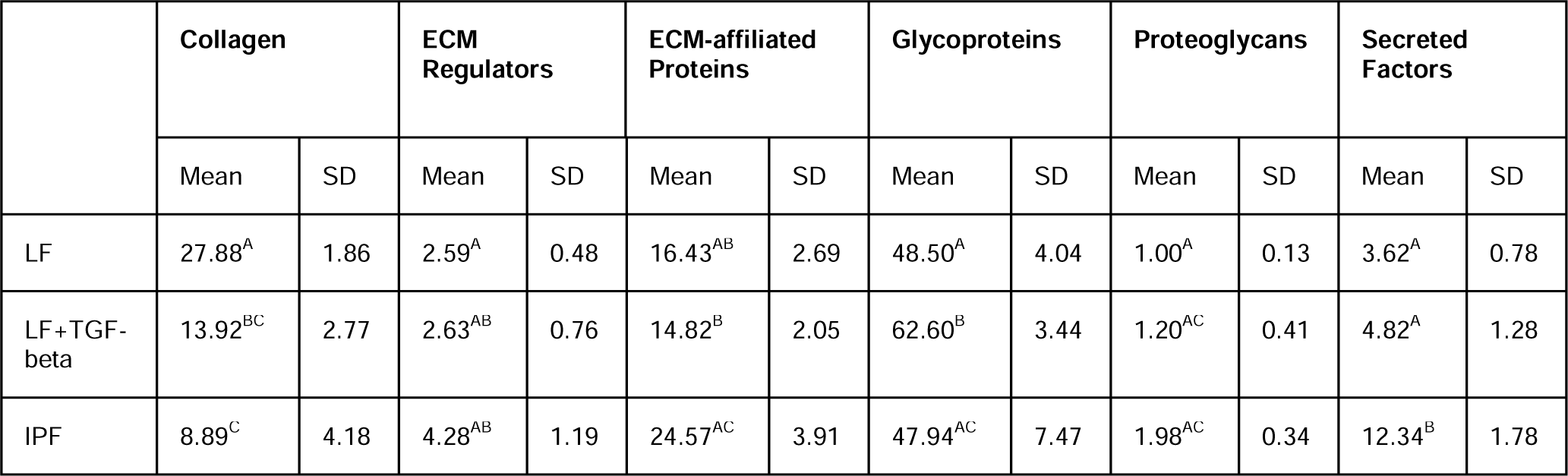
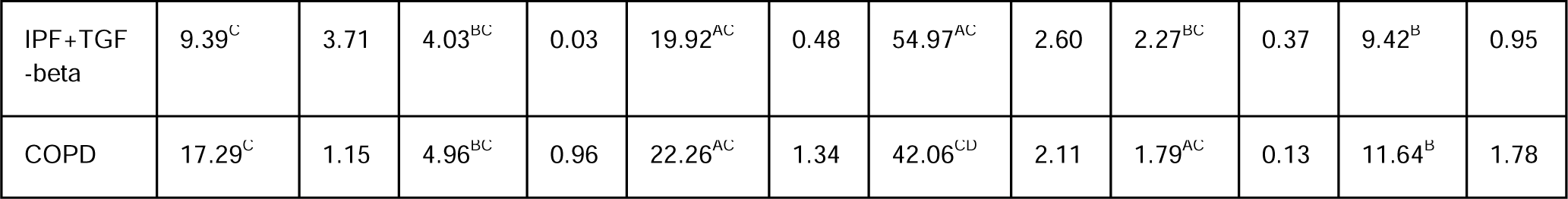
Proteomics analysis of human lung microtissues. One-way ANOVA with post-hoc Tukey HSD test, or their non-parametric equivalent, were used to examine statistical significance between the five human lung microtissues for each of the six ECM protein categories. Values that do not share the same superscript letters are significantly different (p-value ≤ 0.05).

|  | Collagen |  | ECM Regulators |  | ECM-affiliated Proteins |  | Glycoproteins |  | Proteoglycans |  | Secreted Factors |  |
| --- | --- | --- | --- | --- | --- | --- | --- | --- | --- | --- | --- | --- |
|  | Mean | SD | Mean | SD | Mean | SD | Mean | SD | Mean | SD | Mean | SD |
| LF | 27.88<sup>A</sup> | 1.86 | 2.59<sup>A</sup> | 0.48 | 16.43<sup>AB</sup> | 2.69 | 48.50<sup>A</sup> | 4.04 | 1.00<sup>A</sup> | 0.13 | 3.62<sup>A</sup> | 0.78 |
| LF+TGF-beta | 13.92<sup>BC</sup> | 2.77 | 2.63<sup>AB</sup> | 0.76 | 14.82<sup>B</sup> | 2.05 | 62.60<sup>B</sup> | 3.44 | 1.20<sup>AC</sup> | 0.41 | 4.82<sup>A</sup> | 1.28 |
| IPF | 8.89<sup>C</sup> | 4.18 | 4.28<sup>AB</sup> | 1.19 | 24.57<sup>AC</sup> | 3.91 | 47.94<sup>AC</sup> | 7.47 | 1.98<sup>AC</sup> | 0.34 | 12.34<sup>B</sup> | 1.78 |
| IPF+TGF<br>-beta | 9.39<sup>C</sup> | 3.71 | 4.03<sup>BC</sup> | 0.03 | 19.92<sup>AC</sup> | 0.48 | 54.97<sup>AC</sup> | 2.60 | 2.27<sup>BC</sup> | 0.37 | 9.42<sup>B</sup> | 0.95 |
| COPD | 17.29<sup>C</sup> | 1.15 | 4.96<sup>BC</sup> | 0.96 | 22.26<sup>AC</sup> | 1.34 | 42.06<sup>CD</sup> | 2.11 | 1.79<sup>AC</sup> | 0.13 | 11.64<sup>B</sup> | 1.78 |

**Supplementary Table 1** lists all proteins that are either significantly up-regulated or down-regulated in each of the comparisons. Notably, some collagens were down-regulated in IPF, healthy + TGF-beta and COPD as compared to healthy. Collagen triple helix repeat-containing protein 1, a secreted protein found in injured and diseased arteries that was shown to inhibit collagen expression and promotes cell migration, was up-regulated in both IPF and COPD as compared to healthy. Fibronectin was up-regulated in both IPF and healthy + TGF-beta as compared to healthy. Protein-lysine 6-oxidase (involved in protein crosslinking) and procollagen-lysine,2-oxoglutarate 5-dioxygenase 2 (involved in collagen cross-linking), were up-regulated in IPF as compared to healthy. Fibronectin was up-regulated in IPF as compared to COPD.

### Decellularized Lung Microtissues Support Human Breast Cancer Cell Culture

We investigated whether decellularizing lung microtissues to generate cultured lung ECM can support the viability and proliferation of cancer cells, to elucidate whether ECM generated by human cells with different disease states may affect cancer cell growth. Highly metastatic, GFP nuclear-labeled MDA-MB-231 breast adenocarcinoma cells were seeded in low adhesion microwells with or without cultured healthy or cultured IPF lung ECM for 10 days (**Figure 7**). Both healthy and IPF ECM supported greater cell proliferation than in the absence of ECM.

**Figure 7.**
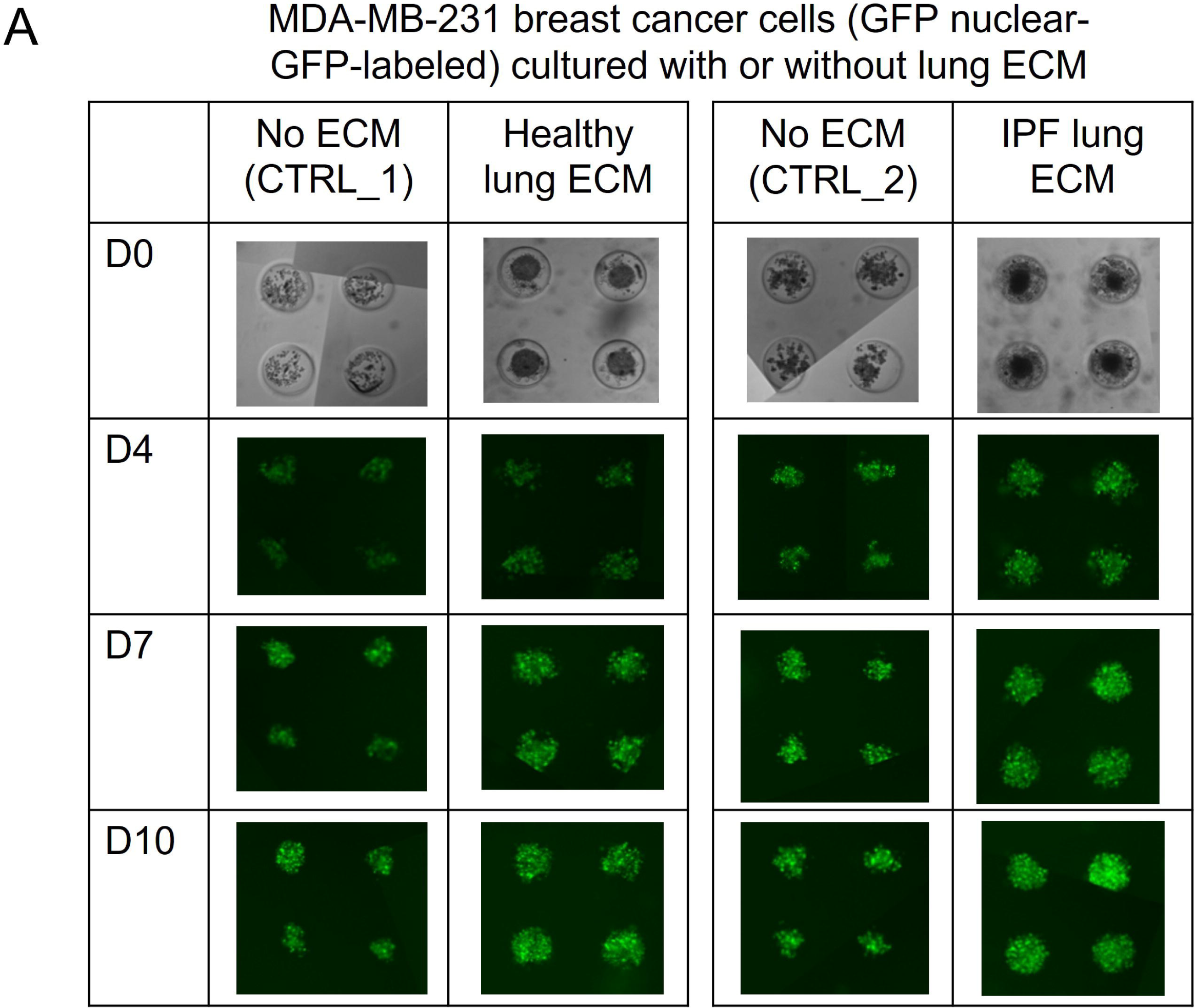
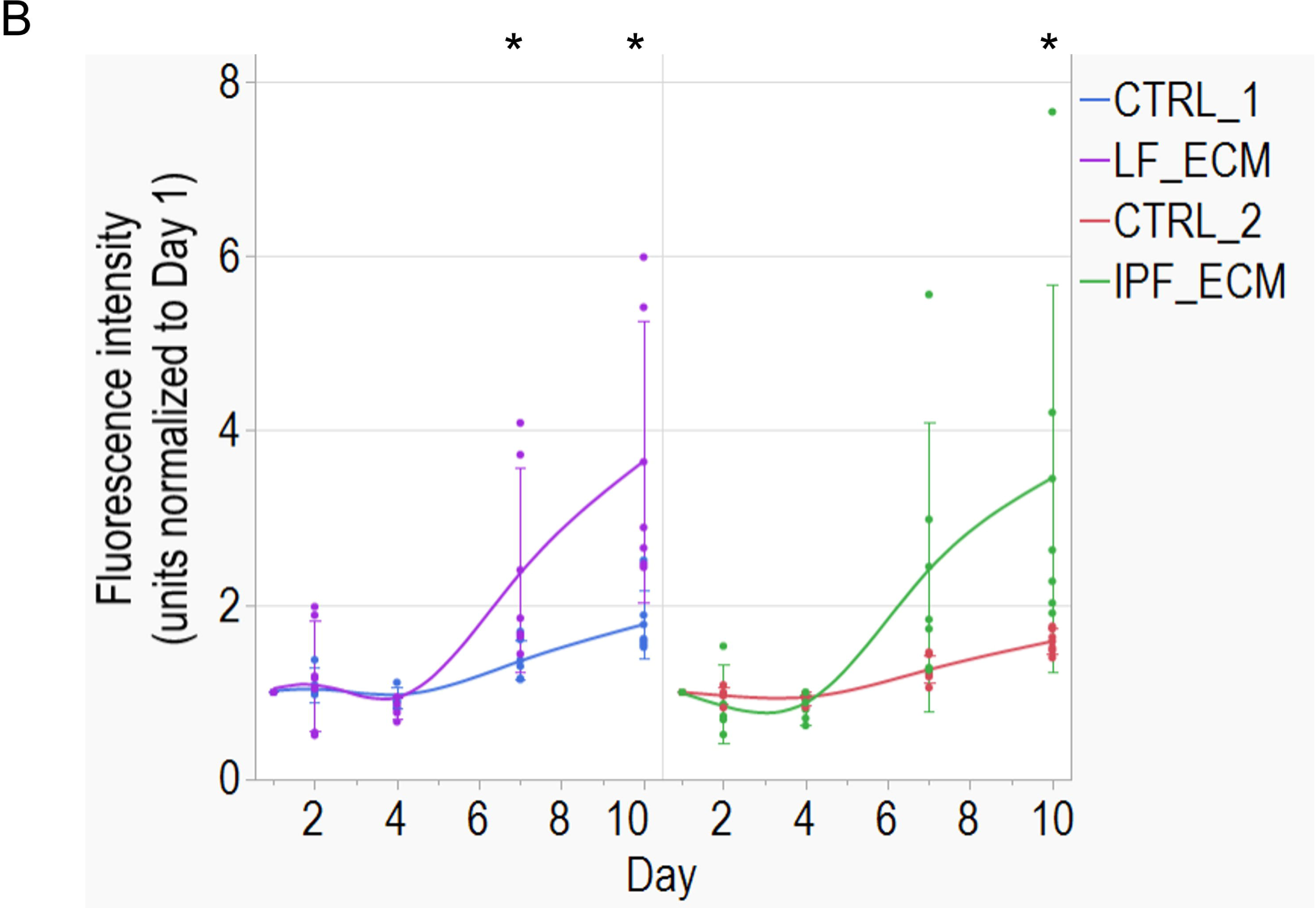
Decellularized lung ECM promoted proliferation of highly metastatic breast cancer cells. Highly metastatic, GFP nuclear-labeled MDA-MB-231 breast adenocarcinoma cells were seeded into decellularized agarose gels with or without cultured healthy or IPF lung ECM for 10-days to evaluate breast cancer cell proliferation with fluorescence microscopy (A). Both healthy and IPF ECM supported greater cell proliferation than in the absence of ECM (B). Student’s *t*-test or the non-parametric equivalent was used to examine statistical significance between with or without ECM at each of the time points. * denotes statistical significance (p-value ≤ 0.05).

### Cultured Cardiac Fibroblast ECM is stiffer than cultured healthy lung ECM

Mechanical characterization using atomic force microscopy (**Figure 8**) showed that cultured cardiac ECM exhibited relatively high stiffness of 35 kPa, relative to lung fibroblast ECM of 4 kPa. This is comparable to in vivo values [40, 41], and higher than typical values of Matrigel and collagen [42], and much lower than stiff plastic culture surfaces of 2 to 4 GPa [40]. Of note, the elastic modulus for cultured IPF lung ECM was marginally stiffer but more variable than cultured healthy lung ECM, which may occur due toto patchy and nodule-like collagen, as observed in SHG (**Figure 6**).

**Figure 8.**
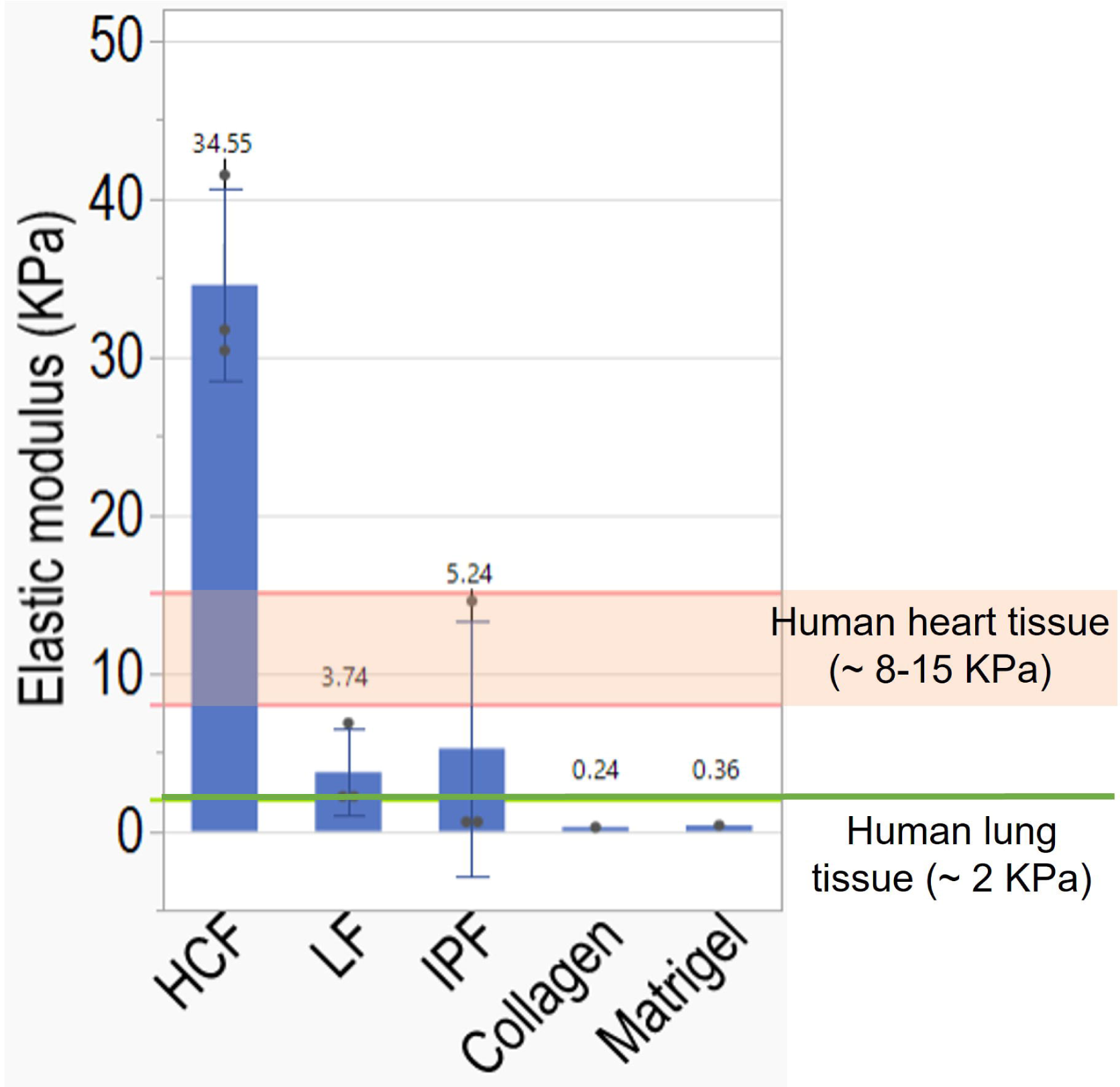
Tissue and disease specific stiffness of decellularized microtissues fabricated with healthy human cardiac (HCF), healthy human lung (LF) or idiopathic fibrotic lung (IPF) fibroblasts. Atomic force microscopy was used to characterize the elastic modulus of three different microtissues immediately after decellularization, with values for collagen [42], Matrigel [42], human heart tissue [40], and human lung tissue [41] for comparison.

### Differential gene expression of healthy cardiac and healthy lung microtissues

We examined the transcriptomics of healthy lung and cardiac microtissues with RNAseq to elucidate the differential gene expression and potential molecular pathways that may be involved in the difference we observed in their cultured ECM. The principal component analysis (PCA; **Figure 9A**) of RNAseq results suggested that most of the variance (99.69%) was explained by the two tissue-specific microtissues, and less than 1% was explained by within group variability. **Figure 9B** reveals the most significant pathways that were tissue-dependent include ECM organization, negative regulation of apoptotic process, positive regulation of cell migration, and angiogenesis. DESeq2 analysis showed that healthy lung microtissues had greater gene expression of collagen genes including COL1A1, COL4A5, COL4A6, COL5A3, COL6A1, COL6A2, COL6A3, COL12A1, which correspond to the differential ECM proteome of the two microtissues (**Supplementary Table 1**).

**Figure 9.**
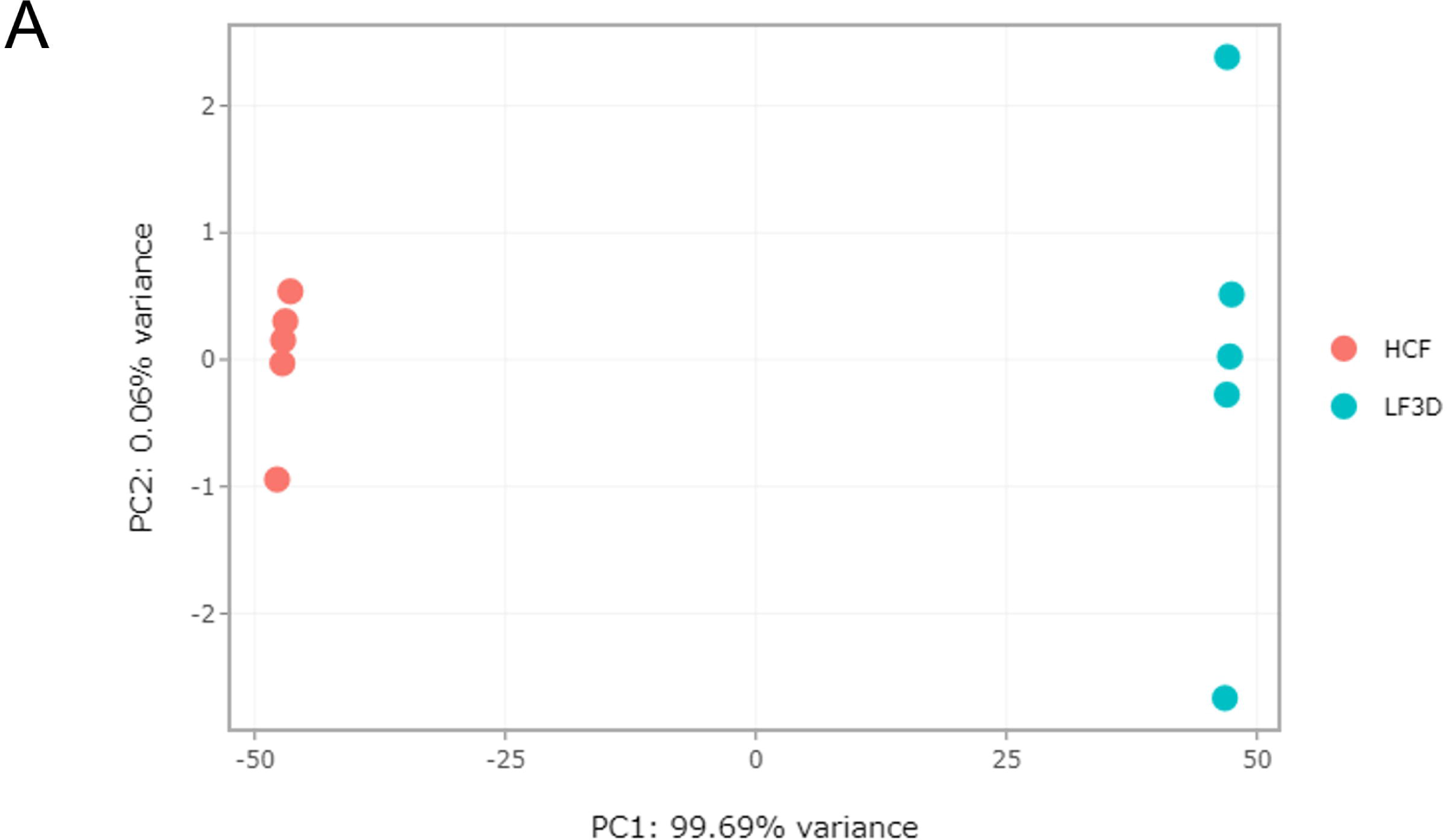
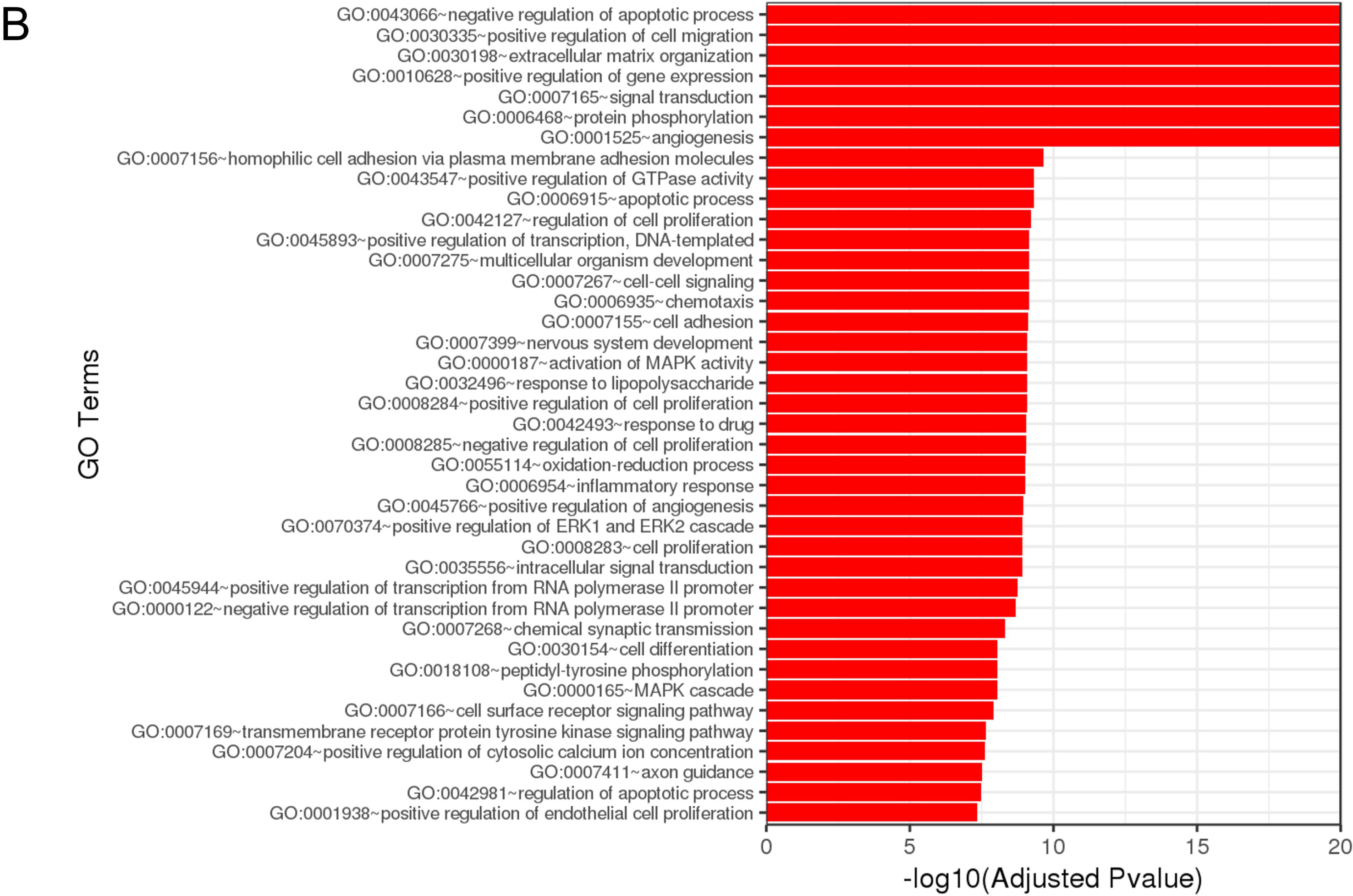
Expression of genes in lung vs heart microtissues. The principal component analysis (A) of RNAseq results suggested that 99.69% of variance was explained tissue specificity of microtissues. The enriched gene ontology terms that were generated by clustering significantly differentially expressed genes using Fisher exact test revealing tissue-specific pathways (B).

## Discussion and Conclusion

Cardiac tissue engineering scaffolds should ideally present a variety of ECM ligands to engage myocytes and endothelial cells and exhibit robust mechanical properties [43]. We show that cell-secreted ECM from human cardiac fibroblasts exhibit protein composition comparable to human and porcine heart tissues, which support the culture of cardiac myocytes and endothelial cells. This represents a promising and scalable approach for regenerative human cardiac ECM, which may minimize immunogenicity in human patients. Further experiments could explore co-culture conditions with other stromal and immune cells to further optimize ECM composition and architecture.

Our comparison of ECM secreted by healthy and fibrotic lung fibroblasts show pronounced differences in microstructure and composition, resulting in enhanced adhesion and proliferation of metastatic breast cancer cells in ECM relative to none. Recent work has elucidated a lung metastatic niche with aberrant ECM composition (e.g. increased collagen and proteolycan deposition) as well as enhanced ECM crosslinking and alignment [44] [45]. This dysregulation is reminiscent of fibrosis associated with chronic lung diseases [46]. Our proteomic analyses show upregulation of proteins involved in protein-crosslinking and promote cell migration in IPF and COPD relative to healthy, analogous to the lung metastatic niche, but also a reduction in collagens in IPF and COPD related to healthy. This results suggest that additional factors beyond fibroblasts (e.g. inflammatory signals) could contribute to the lung metastatic niche.

In conclusion, we demonstrate a method for preparing human fibroblast secreted ECM by decellularizing engineered microtissues in low adhesion microwells. Importantly, cultured ECM exhibited tissue and disease-specific phenotypes (e.g. ECM proteome, mechanical properties, collagen architecture). Overall, this approach enables manufacturing of personalized human ECM specific to the tissue of origin, with the potential for new therapeutic treatments and disease modeling.

## Authors’ Contribution

**VF and VV:** Investigation

**BCI:** Conceptualization, Formal Analysis, Funding Acquisition, Investigation, Methodology, Project Administration, Supervision, Writing

## Acknowledgement

We thank Prof. Eric Darling at Brown University for his advice and support for the mechanical examination of the microtissues, as well as Dr. Geoff Williams for his expertise in multiphoton second harmonic generation microscopy for imaging collagen. We also thank the Brown University Molecular Pathology Core, the Brown University Leduc Bioimaging Facility, and The University of Oklahoma Mass Spectrometry, Proteomics, and Metabolomics Core Facility for their support.

## Supplementary Tables

**Supplementary Table 1:** Differentially regulated proteins between microtissues.

| Comparison<br>(Group 1 vs 2) | Upregulated in Group 1 vs 2 | Downregulated in Group 1 vs 2 |
| --- | --- | --- |
| **HCF vs HCF_M** | Fibronectin, Decorin, Collagen alpha-2(IV) chain, Collagen alpha-1(V) chain, Laminin subunit alpha-2 | EMILIN-2, Vitronectin, Von Willebrand factor |
| **LF vs LF+TGF-beta** | Periostin, laminin subunit beta-2, ADAMTS-like protein 4, Collagen alpha-3(V) chain, Isoform 1 of Collagen alpha-1(II) chain, Collagen alpha-1(XXI) chain, Collagen alpha-5(IV) chain, Prolyl 3-hydroxylase 1 | Fibronectin, biglycan, Isoform 6 of Elastin, Fibulin-5, Isoform 5 of Elastin, collagen alpha-1(VIII) chain, Vitronectin, |
| **LF vs IPF** | Periostin, Isoform 4 of Collagen alpha-3(VI), Collagen alpha-3(VI) chain, Collagen alpha-1(VI) chain, laminin subunit beta-2, Collagen alpha-1(I) chain, Collagen alpha-2(I) chain, Collagen alpha-1(V) chain, Collagen alpha-2(V) chain, Collagen alpha-3(V) chain, Fibulin-5, Collagen alpha-1(XXI) chain, Isoform 1 of Collagen alpha-1(II) chain, Collagen alpha-6(IV) chain, Collagen alpha-5(IV) chain | Collagen triple helix repeat-containing protein 1 (secreted by injured/disease cells), Protein-lysine 6-oxidase (protein crosslinking), procollagen-lysine,2-oxoglutarate 5-dioxygenase 2 (collagen cross-linking), Vitronectin, |
| **LF vs COPD** | Galectin-1, Periostin, Collagen alpha-3(VI) chain, Collagen alpha-1(VI) chain, laminin subunit beta-2, Laminin subunit gamma-1, Collagen alpha-2(VI) chain, Galectin-3, Laminin subunit alpha-4, Isoform 2 of Latent-transforming growth factor beta-binding protein 4, Collagen alpha-3(V) chain, Collagen alpha-6(IV) chain, Collagen alpha-5(IV) chain, Agrin | Isoform C of Fibulin-1, Collagen triple helix repeat-containing protein 1 (secreted protein in injured and diseased arteries, inhibits collagen expression and promotes cell migration), |
| **IPF vs IPF+TGF-beta** | None | Fibulin-5, Procollagen galactosyltransferase 2, Agrin, Dipeptidyl peptidase 4 (associated with lung diseases) |
| **IPF vs COPD** | Galectin-1, Fibronectin, Galectin-3, Isoform 2 of Collagen alpha-1(XVIII) chain, Insulin-like growth factor-binding protein 3, collagen alpha-1(XVI) chain, von Willebrand factor A domain-containing protein 5A, collagen alpha-1(VIII) chain, Collagen alpha-1(XIV) chain, Dipeptidyl peptidase 9, | Isoform 4 of Collagen alpha-3(VI) chain, Fibulin-5, Heme oxygenase 2, Isoform 1 of Collagen alpha-1(II) chain, |
| **LF vs HCF** | Isoform 4 of Collagen alpha-3(VI) chain, Collagen alpha-3(VI) chain, Collagen alpha-2(VI) chain, Collagen alpha-1(VI) chain, Collagen alpha-1(I) chain, Collagen alpha-2(I) chain, Collagen alpha-2(IV) chain, Collagen alpha-1(IV) chain, Collagen alpha-5(IV) chain, collagen alpha-3(IV) chain, Collagen alpha-1(V) chain, Collagen alpha-3(V) chain, Collagen alpha-1(XII) chain, Collagen alpha-1(XV) chain, Periostin, Transforming growth factor-beta-induced protein ig-h3, Procollagen C-endopeptidase enhancer 1, Fibrillin-2, microfibrillar-associated protein 1, Microfibrillar-associated protein 2, ADAMTS-like protein 1, ADAMTS-like protein 4, Latent-transforming growth factor beta-binding protein 3, laminin subunit beta-2, Laminin subunit alpha-5, Galectin-3, | None |

## Supplementary Methods

### LC-MS/MS identification and label free quantitation of ECM proteins

Samples (cell pellets) were lysed with a lysis buffer (8 M urea, 1 mM sodium orthovanadate, 20 mM HEPES, 2.5 mM sodium pyrophosphate, 1 mM β-glycerophosphate, pH 8.0, 20 min, 4°C) followed by sonication at 40% amplification by using a microtip sonicator (QSonica, LLC, Model no. Q55) and cleared by centrifugation (14 000 × g, 15 min, 15°C). Protein concentration was measured (Pierce BCA Protein Assay, Thermo Fisher Scientific, IL, USA) and a total of 100 μg of protein per sample was subjected for trypsin digestion.

In-solution digestion with trypsin/LysC (V5071, Promega) was performed according to the manufacturer protocol. Tryptic peptides were desalted using C18 Sep-Pak plus cartridges (Waters, Milford, MA) and were lyophilized for 12 hours to dryness. The dried peptides were reconstituted with 200 μl buffer A (0.1 % formic acid). The final concentration of the peptide samples 0.5 μg/μl and 4 μl (2 μg)/sample was injected for LC-MS/MS analysis. The LC-MS/MS was performed on a fully automated proteomic technology platform that includes a Dionex UltiMate ® 3000 (Thermo Fisher Scientific, USA) system connected to a Q Exactive HF-X mass spectrometer (Thermo Fisher Scientific, Waltham, MA). The MS/MS analysis was performed according to the earlier published protocol (Bishop et al., 2021,bioRxiv, doi: https://doi.org/10.1101/2021.02.05.429978).

Peptide spectrum matching of MS/MS spectra of each file was searched against the UniProt Human database (TaxID:9606) using the Sequest algorithm within Proteome Discoverer v 2.4 software (Thermo Fisher Scientific, San Jose, CA). The Sequest database search was performed with the following parameters: trypsin enzyme cleavage specificity, 2 possible missed cleavages, 10 ppm mass tolerance for precursor ions, 0.02 Da mass tolerance for fragment ions. Search parameters permitted dynamic modification of methionine oxidation (+15.9949 Da) and static modification of carbamidomethylation (+57.0215 Da) on cysteine. Peptide assignments from the database search were filtered down to a 1% FDR. The relative label-free quantitative and comparative among the samples were performed using the Minora algorithm and the adjoining bioinformatics tools of the Proteome Discoverer 2.4 software.

